# Optical photothermal infrared spectroscopy can differentiate equine osteoarthritic plasma extracellular vesicles from healthy controls

**DOI:** 10.1101/2022.03.11.483922

**Authors:** Emily J Clarke, Cassio Lima, James R Anderson, Catarina Castanheira, Alison Beckett, Victoria James, Jacob Hyett, Roy Goodacre, Mandy J Peffers

## Abstract

**Background:** Equine osteoarthritis is a chronic degenerative disease of the articular joint, characterised by cartilage degradation resulting in pain and reduced mobility and thus is a prominent equine welfare concern. Diagnosis is usually at a late stage through radiographic examination, whilst treatment is symptomatic not curative. Extracellular vesicles are small nanoparticles that are involved in intercellular communication. The objective of this study was to investigate the feasibility of Raman and optical photothermal infrared spectroscopy to detect osteoarthritis using plasma-derived extracellular vesicles.

**Methods:** Plasma samples were derived from thoroughbred racehorses. A total of 14 samples were selected (control; n= 6 and diseased; n=8). Extracellular vesicles were isolated using differential ultracentrifugation and characterised using nanoparticle tracking analysis, transmission electron microscopy, and human tetraspanin chips. Samples were then analysed using Raman and optical photothermal infrared spectroscopy.

**Results:** Infrared spectra were analysed between 950-1800 cm^-1^. Raman spectra had bands between the wavelengths of 900-1800 cm^-1^ analysed. Bands below 900 cm^-1^. Spectral data for both Raman and optical photothermal infrared spectroscopy was used to obtain a classification model and confusion matrices, characterising the techniques ability to distinguish diseased samples. Optical photothermal infrared spectroscopy could differentiate osteoarthritic extracellular vesicles from healthy with good classification (93.4%) whereas Raman displayed poor classification (64.3%). Plasma-derived extracellular vesicles from osteoarthritic horses contained increased signal for proteins, lipids and nucleic acids.

**Discussion/ conclusion:** For the first time we demonstrated the ability to use optical photothermal infrared spectroscopy to interrogate extracellular vesicles and osteoarthritis-related samples. Optical photothermal infrared spectroscopy was superior to Raman in this study, and could distinguish osteoarthritis samples, suggestive of its potential use diagnostically to identify osteoarthritis in equine patients. This study demonstrates the potential of Raman and optical photothermal infrared spectroscopy to be used as a diagnostic tool in clinical practice, with the capacity to detect changes in extracellular vesicles from clinically derived samples.

## Introduction

Osteoarthritis (OA) is a common degenerative disease of the synovial joint, characterised by catabolic processes observed in articular cartilage, and a notable imbalance in bone remodelling. It results in pain, inflammation and reduced mobility (Mustonen and Nieminen, 2021). OA is the most prevalent cause of equine lameness, with over 60% of horses developing OA within their lifetime; a significant welfare concern (McIlwraith *et al*, 2012). It is a complex heterogeneous condition of multiple causative factors, including mechanical, genetic, metabolic and inflammatory pathway involvement (Clarke *et al*, 2021). OA pathophysiology is conserved across species, resulting in synovitis, cartilage degradation, osteophyte formation, subchondral bone sclerosis, fibrosis and reduced elastoviscosity of synovial fluid found within the joint capsule (Kuyinu *et al*, 2016). The horse is both of interest as a target species for veterinary equine medicine as well as a model organism to study osteoarthritis due to these established similarities, as well as comparable anatomic structures of the human carpal joint with equine carpal and metacarpophalangeal joint (McIlwraith *et al*, 2012).

Extracellular vesicles (EVs) are nanoparticles enveloped in a phospholipid bilayer membrane, secreted by most mammalian cells, that transport biologically active cargo, such as proteins, RNAs, DNAs, lipids, and metabolites (Boere *et al*, 2018). EVs are divided into subgroups as determined by their size and biogenesis. EVs elicit their effects through paracrine signalling, proving fundamental in intercellular communication (Herrmann *et al*, 2021). EVs have been implicated in the propagation of OA, and have been shown to be released and enter chondrocytes, synoviocytes and inflammatory cells (Esa *et al*, 2019; Withrow *et al;* 2016 and Lin *et al*, 2021). Interestingly EVs can serve as disease propogators; promoting an increased expression of cytokines, chemokines and matrix degrading proteinases, or disease preventing; increasing cellular differentiation and reducing apoptosis.

Vibrational spectroscopic methods, including Raman and infrared spectroscopies, provide molecular information about the main molecular constituents commonly found in biological samples such as proteins, lipids, nucleic acids, and carbohydrates based on bond-specific chemical signatures in a non-invasive, non-destructive, and label-free manner (Paraskevaidi *et al*, 2021). In Raman spectroscopy, photons from a monochromatic source interact with the sample and a small fraction of them are inelastically scattered with either higher or lower energies compared with the excitation wavelength. The energy difference between incident and scattered photons corresponds to a Raman shift and it is associated with the chemical structure of molecules in the sample (Lima *et al*, 2021). Raman spectroscopy can discriminate between cell and tissue types and detect chemical alterations prior to morphological changes in various pathological states. It has previously been used to assess the purity of EV preparations (Gualerzi et al, 2019), as well as identify cellular origin of mesenchymal stem cell (MSC)-derived EVs (Gualerzi et al, 2017). Infrared spectroscopy is based on the absorption of infrared radiation by molecular vibrations from bonds that possess an electric dipole moment that can change by atomic displacement (Baker *et al*, 2014).

Although Raman and infrared spectroscopies provide molecular information about the overall biochemistry of samples by monitoring the internal motion of atoms in molecules, each method has advantages and disadvantages for different types of samples due to their different working principles. Thus, having both techniques combined in one single technology is a powerful tool with promising applications. More recently, a novel far-field optical technique has been developed in order to acquire Raman and infrared signatures simultaneously. In this method, the infrared signatures are collected via optical photothermal infrared spectroscopy (O-PTIR), which is based on a pump-probe configuration that couples a tuneable infrared quantum cascade laser (QCL) acting as pump and a visible laser to probe the thermal expansion resulting from the temperature rise induced by the QCL. The probe laser also acts as excitation source for acquiring Raman spectrum simultaneously with infrared data at the same spatial resolution. This scheme has been used to interrogate tissue samples, mammalian cells (Banas *et al*, 2021; Spadea *et al*, 2021) and bacteria (Lima *et al*, 2021) but this is the first study, to our knowledge, to use it in EV or OA work.

We hypothesised that Raman and O-PTIR can be used to identify potential biomarkers of OA using plasma-derived EVs.

## Materials and Methods

### Sample Selection

Plasma samples were collected in accordance with the Hong Kong Jockey Club owner consent regulations (VREC561). All samples were from thoroughbred racehorses, with the donor cohort having a mean age (+/-SEM) of 6.57 +/- 0.45. Horses were selected based on histological scoring of OA severity using a modified Mankin score (McIlwraith *et al*, 2010). A total of 14 samples were selected (control; n= 6 (mean score +/- SEM = 1.83 +/-0.48) and diseased; n=8 (mean score +/- SEM= 16.25 +/- 1.15).

### Extracellular Vesicle Isolation – Differential Ultracentrifugation

Equine plasma samples underwent differential ultracentrifugation (dUC) in order to isolate EVs. Samples were subjected to a 300g spin for 10 minutes, 2000g spin for 10 minutes, 10,000g spin for 30 minutes in a bench top centrifuge. Samples were then transferred to Beckman Coulter thick wall polycarbonate 4ml ultracentrifugation tubes, and spun at 100,000g for 70 minutes at 4°C (Optima XPN-80 Ultracentrifuge, Beckman Coulter, California, USA) in a 45ti fixed angle rotor, with the use of a 13mm diameter Delrin adaptor. Sample pellets were suspended in 50μl of filtered phosphate buffered saline (PBS) (Gibco™ PBS, pH 7.4 - Fisher Scientific, Massachusetts, USA), resulting in 50μl plasma EV (P-EV) samples.

### Extracellular Vesicle Characterisation

#### Nanoparticle Tracking Analysis

Nanoparticle tracking analysis (NTA) was used to quantify EV concentration and size of all samples, using a NanoSight NS300 (Malvern, UK). All samples were diluted in filtered PBS 1:50 (10μl of sample used), to a final volume of 500μl. For each measurement, three 1-min videos were captured, (at a screen gain of 4 and detection threshold of 12. After capture, the videos were analysed by the in-build NanoSight Software NTA 3.1 Build 3.1.46. Hardware: embedded laser: 45 mW at 488 nm; camera: sCMOS.

#### Transmission Electron Microscopy

EV presence and morphology were characterised using transmission electron microscopy. 10 μl of each sample was placed onto a carbon coated glow discharged grid and incubated at room temperature for 20 minutes. Samples were then subject to a negative staining protocol. EVs were fixed onto the grid with 1% glutaraldehyde for 5 minutes. The sample grids were incubated on 1% aqueous uranyl acetate (UA) (Thermofisher Scientific, Massachusetts, USA), for 60 seconds, followed by 4%UA/2% Methyl Cellulose (Sigma Aldrich, Gillingham, UK) at a 1:9 ratio on ice for 10 minutes. Grids were then removed with a 5mm wire loop and dried. The prepared grids were then viewed at 120KV on a FEI Tecnai G2 Spirit with Gatan RIO16 digital camera.

#### Exoview Characterisation

The exoview platform (NanoView Biosciences, Malvern Hills Science Park, Malvern) was used to determine EV concentration, surface marker identification and to perform fluorescent microscopy and tetraspanin colocalization analysis. We had previously tested both the human and murine chips on equine samples and demonstrated the human chips were more compatible (data not shown). ExoView analyses EVs using visible light interference for size measurements and fluorescence for protein profiling. Samples were analysed in triplicate using the ExoView Tetraspanin Kit (NanoView Biosciences, USA) and were incubated on the human ExoView Tetraspanin Microarray Chip for 16 hours at room temperature. Following this sample chips were incubated with tetraspanin labelling antibodies, namely anti-CD9 CF488, anti-CD81 CF555 and anti-CD63 CF647 and the MIgG negative control. The antibodies were diluted 1:500 in PBST with 2% BSA. The chips were incubated with 250 μL of the labelling solution for 1 hour. The sample chips were washed and imaged with the ExoView R100 reader ExoView Scanner v3.0. Data was analysed using ExoView Analyzer v3.0. Fluorescent cut offs were set relative to the MIgG control. Total EVs were determined as the number of detected particles bound to tetraspanin antibodies (CD9, CD81, CD63) and normalised to MIgG antibody.

#### Raman Spectroscopy and Infrared Spectroscopy (O-PTIR)

For all samples O-PTIR measurements were acquired on single-point mode using a mIRage infrared microscope (Photothermal Spectroscopy Corp., Santa Barbara, USA), with the pump consisting of a tuneable four-stage QCL device, while the probe beam is a continuous wave (CW) 532 nm laser. Spectral data were collected in reflection mode using a 40×, 0.78 NA, and 8 mm working distance Schwarzschild objective. Single-point spectral data were acquired over a spectral region of 930-1800 cm^-1^, with 2cm^-1^ spectral resolution and 10 scans per spectrum. Raman data were acquired using a Horiba Scientific iHR-320 spectrometer coupled to mIRage, using a grating of 600 l/mm, 10 s as acquisition time, spectral region of 500-3400 cm^-1^, with 2 cm^-1^ spectral resolution and 10 scans per spectrum.

#### Statistical Analysis

Raman and O-PTIR spectroscopic data was analysed using principal component analysis (PCA) to determine each techniques ability to identify diseased samples from healthy. Spectral data was also used as input for partial least squares discriminant analysis (PLS-DA) in order to generate a computerised model. Further analysis involved using a classification model and confusion matrices, whereby bootstrapping was performed 10000 times to permute whether the EV spectrum was classified as OA or control.

## Results

### EV Characterisation

#### Nanoparticle tracking analysis

Particle size and concentration characterisation was performed using NTA. NTA determined the average plasma sample concentration to be 2.02×10^9^ particles/ml. Analysis was suggestive of a heterogeneous population of EVs, ranging from exosomes to microvesicles (Figure 1A).

**Figure 1.**
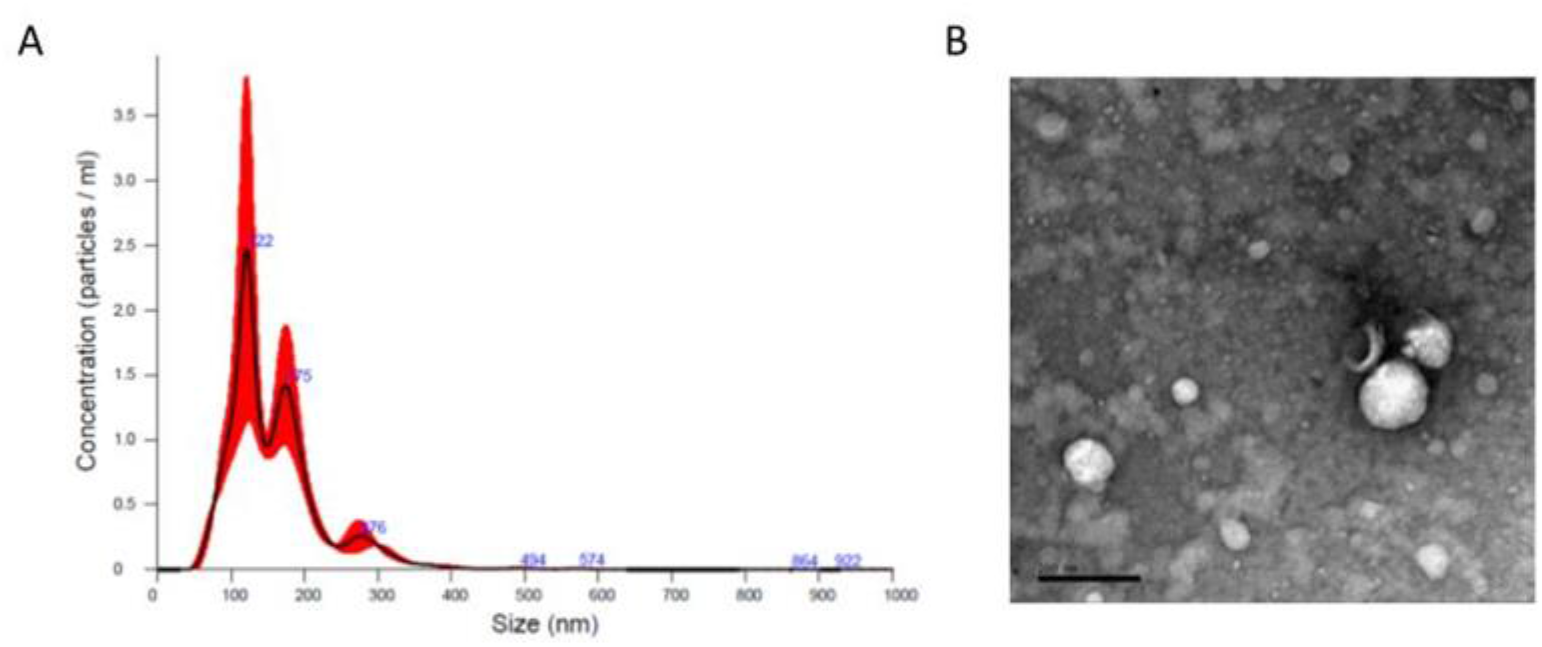
Characterisation of EVs using Nanoparticle Tracking. 1A) Nanoparticle tracking analysis (NTA) of a representative sample of plasma derived EVs. Concentrations of EVs in particles/ml and particle size measured in nm, all measurements recorded using NanoSight NS300, and data analysed by the in-build NanoSight Software NTA 3.1 Build 3.1.46. Hardware: embedded laser: 45 mW at 488 nm; camera: sCMOS. 1B) Transmission electron microscopy (TEM) micrograph of negatively stained representative P-EV samples. Samples fixed to grids were visualised using a FEI Tecnai G2 Spirit with Gatan RIO16 digital camera.

#### Transmission electron microscopy

To confirm the particles isolated from plasma samples were indeed EVs we negatively stained and visualised them using transmission electron microscopy. Spherical structures within EV size ranges (30nm-100nm (exosomes) and 100nm-1000nm (microvesicles)) were identified with a clearly defined peripheral membrane as shown in Figure 1B.

#### Exoview

Exoview was used on a representative pool of plasma samples. The EVs extracted from plasma had the highest particle counts on the CD9 capture spots, equating to a concentration of around 1.4×10^8^ CD9 positive particles/ml. CD81 (7×10^7^ particles/ml) and CD63 (6×10^7^ particles/ml) positive particles were also detectable (Figure 2A). It was observed that most EVs detected were less than 100nm (Figure 2B). It was also found that with plasma EV samples, the greater the expression of CD9 the greater the expression of CD81, whereby a distinct positive correlation can be observed (Figure 2C). Co-localisation analysis was also performed and identifying that 91% of plasma-derived EVs were positive for the CD9 surface tetraspanin, followed by 5% expressing CD81, and 3% expressing both CD81 and CD9. CD63 expression was lowest at 0.8% (Figure 2D). Finally, plasma EVs were visualised using fluorescent microscopy, as shown in Figure 2E.

**Figure 2.**
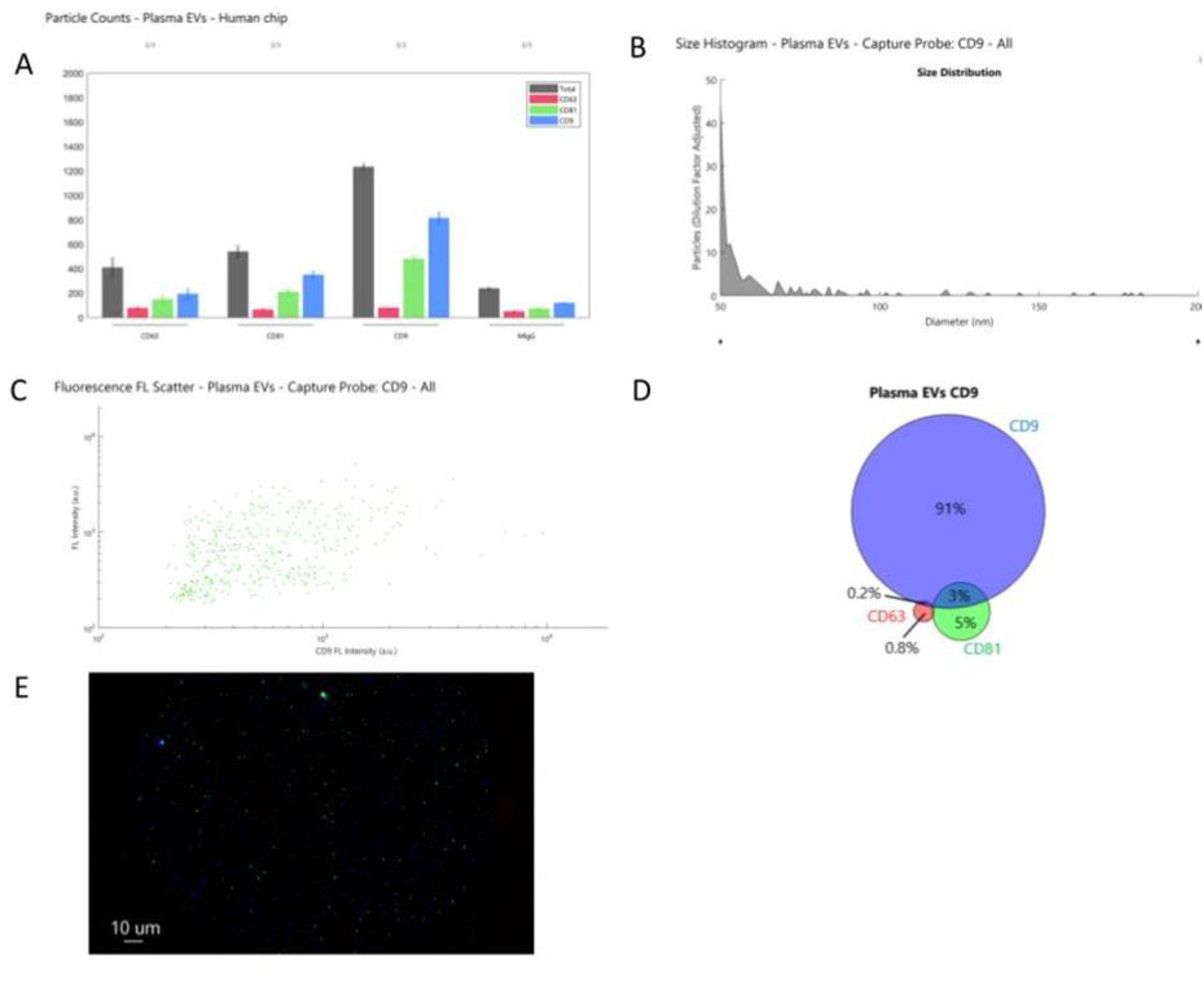
Characterisation of EVs using Exoview. A) Particle counts for plasma EVs from the human tetraspanin chip. B) A size histogram of plasma EV samples as captured on the CD9 human tetraspanin chip. C) Scatter diagram demonstrating correlation between the number of CD9 positive EVs (X axis) and CD81 positive EVs (Y axis). D) colocalization analysis of the presence of surface tetraspanins on equine plasma EVs. E) fluorescent microscopy visualising plasma-derived EVs, with colour denoting surface tetraspanin positive identification (red – CD63, blue- CD9, and green -CD81).

#### Raman Spectroscopy and Infrared Spectroscopy (O-PTIR)

Infrared and Raman spectral data collected from healthy and diseased plasma-derived EV samples displayed similar biochemical features with subtle changes in signal intensity, while appearance or disappearance of peaks were not observed (Figure 3A and B).

**Figure 3.**
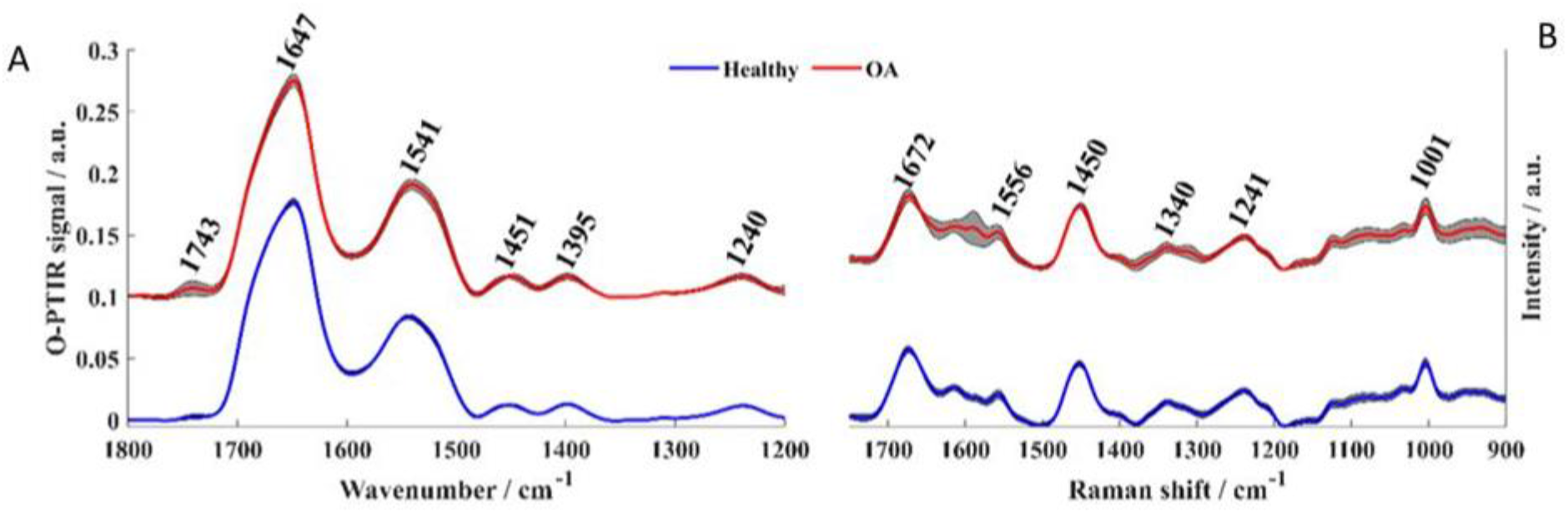
Fingerprint region of averaged spectra. A) Infrared and B) Raman spectra collected from healthy (blue line) and diseased (red line). Plots are offset for clarity.

Infrared signatures acquired by O-PTIR spectroscopy were recorded from 950-1800 cm^-1^, however, bands peaking below 1200 cm^-1^ were removed from the analysis due to interference from bands attributed to minerals from PBS (Mihaly *et al*, 2017). The band peaking at 1743 cm^-1^ arose from C=O ester groups from lipids including phospholipids, triglycerides and cholesterol (Mihaly *et al*, 2017) Amide I vibration, peaking at 1647 cm^-1^ was associated mainly to C=O stretching vibration from peptide bonds in proteins (Mihaly *et al*, 2017; Barth,2007; Paolini *et al*, 2020). Amide II band absorption was found in 1544 cm^-1^ and is attributed to the out-of-phase combination of the N-H in-plane bend and C-N stretching vibration with smaller contributions from the C-O in-plane bend and the C-C and N-C stretching vibrations of peptide groups (Mihaly *et al*, 2017; Barth, 2007; Paolini *et al*, 2020). The band observed at 1240 cm^-1^ results from the coupling between C-N and N-H stretching from proteins (amide III) (Barth, 2007), but it was also influenced by PO2^-^ asymmetric stretching from phosphodiester bonds in nucleic acids (Paraskevaidi *et al*, 2021). The peak at 1451 cm^-1^ corresponded to bending vibration (scissoring) of acyl CH_2_ groups in lipids (Mihaly *et al*, 2017; Barth et al, 2007; Paolini et al, 2020), whereas the band peaking at 1395 cm^-1^ aroses from COO^-^ symmetric stretching from amino acid side chains and fatty acids (Mihaly *et al*, 2017; Barth, 2007; Paolini *et al*, 2020). Spectral signatures from minerals were also observed in Raman spectrum acquired from healthy and diseased samples in the low wavenumber region (below 900 cm^-1^), therefore, only bands peaking between 900-1800 cm^-1^ were analysed. In Raman spectra, peaks associated to amide I, II, and III from peptide bonds were observed peaking at 1672, 1556, and 1241 cm^-1^ respectively (Zhang *et al* 2020; Gualerzi *et al*, 2019). The band peaking at 1450 cm^-1^ originated from CH_2_/CH_3_ bending vibrations from lipids and proteins, while the peak at 1004 cm^-1^ was attributed to the phenylalanine ring breathing, and the peak at 1340 cm^-1^ was associated to nucleic acids (Zhang *et al* 2020; Gualerzi *et al*, 2019).

Infrared and Raman were subjected to principal component analysis (PCA) in order to examine the ability of both techniques to discriminate healthy and diseased plasma-derived EV samples (Figure 4). PC scores plot obtained from O-PTIR (infrared signatures) as input data showed satisfactory discrimination between healthy and diseased samples (Figure 4A), with scores from control samples grouping on the negative side of PC-1 axis while scores related to OA clustered on positive side of the PC-1 axis. The loadings plot (Figure 4B) revealed positive loadings to all bands displayed in Figure 4a, indicating higher amount of the molecular constituents associated with these vibrations, i.e. proteins, lipids and nucleic acids, in samples derived from plasma EVs in OA. The scores plot obtained by subjecting Raman signatures to PCA displayed poor discrimination between healthy and OA samples (Figure 4).

**Figure 4.**
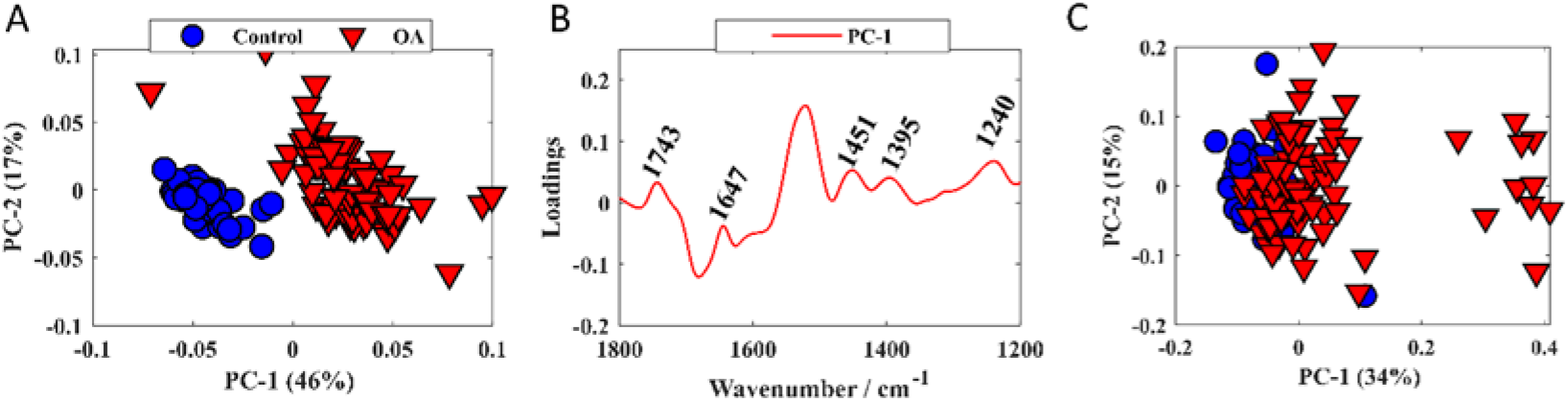
PCA scores and loadings plots. A) PCA score plot from infrared data. B) PCA loadings plot from infrared data. C) PC scores plot of Raman data; values in parentheses are the percentage total explained variance.

Raman and O-PTIR spectral data were used as input for PLS-DA in order to generate a model for classifying OA and control samples based on EV composition. The classification models and confusion matrices obtained by PLS-DA using O-PTIR and Raman data are shown in Figure 5. Bootstrapping was performed 10000 times to permute whether the EV spectrum was classified as OA or control. PLS-DA model obtained from O-PTIR data demonstrated good correct classification rates (CCR) of an average of 93.41%, while PLS-DA model generated by using Raman signatures as input data presented a poor CCR (64.63%). These findings agree with the results obtained by PCA, indicating superior ability of O-PTIR to discriminate healthy from OA samples.

**Figure 5.**
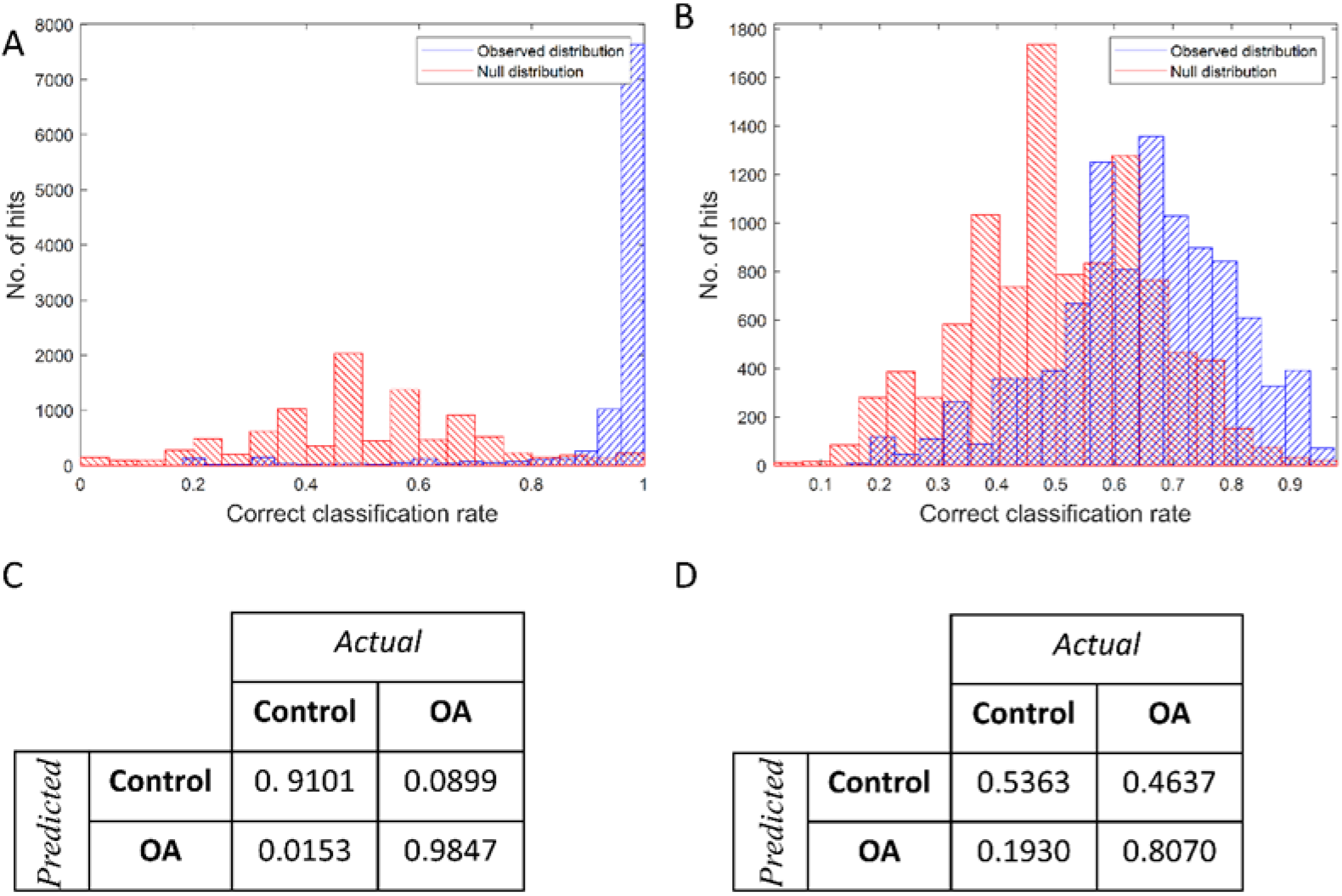
Classification model for PLS-DA of infrared and Raman spectra. A) Infrared and B) Raman spectra showing classification rates for real (blue) and random (red), with a correct classification rate of 93.41% for infrared and 64.63%. C) and D) are the respective average confusion matrices with rows representing predicted classification and columns representing experimental.

## Discussion

This study, for the first time investigated whether O-PTIR could be used successfully to interrogate plasma-derived EVs in control and OA equine samples. We demonstrated that indeed the novel spectroscopy technique O-PTIR was able to determine differences in EV content attributed to disease pathology. Thus O-PTIR could serve as a platform for as future biomarker studies in equine OA. Plasma EV concentration (2.02×10^9^ particles/ml) was quantified using NTA, an accepted method of EV evaluation known for its repeatable and reproducible (Vestad *et al*, 2017). The representative plasma sample shown reflect the biological sample heterogeneity, showing a range of EV sizes indicative of both exosome and microvesicles subgroups. The concentration determined was similar to those reported for plasma EVs in other species within literature. For example, human plasma EVs identified by Palviainen *et al* (2020) 2.46×10^9^-1.10×10^10^ particles per ml. However, a study analysing plasma EVs across the duration of equine endurance racing found that baseline plasma EV concentration was 5.6×10^12^ particle per ml (de Oliveira Jr *et al*, 2021). Exoview analysis identified EVs that were positive for the tetraspanins CD81 and CD9, however a low percentage of EVs were positive for the surface marker CD63. This may a result of poor protein homology between equine and humans. Although, a recent paper using intracellular trafficking demonstrated that CD63 expression in HeLa cells was specific to exosomes, and often a lack of CD63 expression may be due to small microvesicle production, referred to as ectosomes, and that this type of EV is far more prolific than CD63 positive exosomes (Mathieu *et al*, 2021).

Infrared spectroscopy techniques have been used to probe EVs previously in order to identify structural components (Kim *et al*, 2019) as well as proteins, lipid and nucleic acid components, as found in our study (Kim *et al*, 2019). This is the first paper to our knowledge to probe OA EVs using Raman spectroscopy and O-PTIR spectroscopy. In other work, Zhai *et al* (2019) found that in bone, mineral and carbonate content varied significantly with OA stage, with carbonate increasing with OA. They also identified using Fourier transform infrared spectroscopy that acid phosphate, collagen maturity and crystallinity varied with OA. In addition, the use of infra-red spectroscopy is compounded by the findings of Afra *et al* (2017) in a study utilising an experimental model of OA in rats. Here the spectral differences between control and OA samples could be correlated to Mankin score and glycosaminoglycan content. A previous study by our group used attenuated total reflection Fourier-transform infrared (ATR-FTIR) spectroscopy on OA equine serum. This infrared spectroscopy study found separation between groups with 100% sensitivity and specificity, with the six most significant peaks between groups being attributed to proteins and lipids. Similarly, this was observed within our study, with increased abundance found within our OA group. The stated study postulated these observations may be associated to increased lipid and protein expression including increased expression of type 1 collagen, and decreased expression of type 2 collagen characteristic of OA (Paraskevaidi *et al*, 2020).

Previously, changes in plasma and lipid concentration in plasma and serum derived from OA patients has been described. One study utilised serum samples from horses to discriminate proteomic changes due to exercise or the development of early OA (Frisbie *et al*, 2008). Researchers identified six biomarkers with the ability to discriminate OA from exercise groups. For example, the concentration of serum C1,2C (reflective of type 1 and 2 collagen degradation fragments) and collagen 1 was found to significantly increase in OA groups compared with exercise alone (Frisbie *et al*, 2008). In addition, a multiplexed proteomic study on human OA serum identified a panel of 14 candidate biomarkers for OA (Fernandez-Puente *et al*, 2017), utilising cartilage, synovial fluid, chondrocytes and serum. These prospective biomarkers include von Willebrand factor (inflammation and haemostasis) and haptoglobin (an inflammation inducible plasma protein) (Fernandez-Puente *et al*, 2017).

Furthermore, a previous study in plasma identified different lipid profiles in OA using a destabilisation of the medial meniscus model (Pousinis *et al* 2020). Altered lipids included classes of cholesterol esters, fatty acids, phosphatidylcholines, N-acylethanolamines, and sphingomyelins and some were attributed to cartilage degradation.

Raman spectroscopy has been used to characterise EVs and their composition. It has been used to interrogate EVs undergoing autophagy (Chalapathi *et al*, 202), as well as distinguishing EVs derived from bovine placenta and mononuclear cells (Zhang *et al*, 2020), identifying differential features in Parkinson’s disease pathology (Gualerzi *et al*, 2019) and sporadic Amyotrophic Lateral Sclerosis (Morasso *et al*, 2020). Additionally, it has previously been used to successfully discriminate between healthy and diseased joint tissues in order to identify subtle molecular and biochemical changes as a result of disease. Buchwald *et al* (2017) utilised Raman spectroscopy to identify compositional and structural changes in bone from the hip joints of OA patients demonstrating that subchondral bone from OA patients was less mineralised due to a decrease in hydroxyapatite. Furthermore, a study performed by de Souza *et al* (2014) using two *in vivo* experimental rat models of knee OA (treadmill exercise induced and collagenase induced) established molecular signatures unique to OA. Raman ratios relating to mineralization and tissue remodelling were significantly higher in OA groups. Specifically, the ratio between phosphate and amide III has been shown to reflect the degree of mineralisation and carbonate/ amide III is indicative of bone remodelling. De Souza *et al* (2014) also commented on the lack of literature available with regard to the use of Raman spectroscopy for OA research and the importance of this developing field. More recently, Hosu *et al* (2019) reviewed the importance of Raman spectroscopy in identifying pathologically associated crystals such as monosodium urate and calcium pyrophosphate dihydrate in rheumatoid diseases.

We recognise a number of limitations in our study. We were restrained by the number of samples available to us, and our findings need validation in a larger cohort. In addition, large sample volumes were necessary to have an adequate number of EVs for analysis, providing an appropriate signal to noise ratio. In our future studies minimum sample volume required will be optimised. Additionally, we used a single time point ‘snap shot’ of disease. Further work is needed to determine if O-PTIR is sensitive enough to determine differences in a range of OA phenotypes and severities, and correlate differences to specific biological functions of EVs.

Overall this study demonstrates the potential of Raman and OPTIR spectroscopy to be used as a diagnostic tool in clinical practice. Further work is required to identify if OA-related changes in plasma-derived EVs is related to the pathogenesis in the joint. We are currently quantifying the EV cargo using NMR metabolomics, mass spectrometry proteomics and sequencing platforms in order to provide complete characterisation of EVs in OA and determine their contribution to disease propagation.

### Conclusion

In conclusion, EVs derived from equine plasma in OA were probed using Raman and O-PTIR spectroscopy. O-PTIR spectroscopic data was found to be superior in classifying samples from OA patients compared to Raman spectroscopy. O-PTIR spectroscopy is an exciting platform for bedside analysis of plasma to diagnose OA.

## Acknowledgements

Emily Clarke is a self-funded PhD student. Mandy Peffers is funded through a Wellcome Trust Intermediate Clinical Fellowship (107471/Z/15/Z). Our work is also supported by the Medical Research Council (MRC) and Versus Arthritis as part of the MRC Versus Arthritis Centre for Integrated research into Musculoskeletal Ageing (CIMA).

